# Differential Effects of Koisio Technology-Modulated Solutions on the Growth of Lung Fibroblast Cell Cultures and Lung Cancer Cell Cultures

**DOI:** 10.1101/2022.03.02.482600

**Authors:** Mingchao Zhang, Yinghui Men, Qi Zhu, Weihai Ying

## Abstract

Cell growth is a crucial biological property of cells, which plays key roles in several major biological processes including organ development, tissue repair and cancer development. It is of both scientific and medical significance to discover new strategies to modulate cell growth. Koisio technology is a novel technology that modulates the properties of water solely by physical approaches without additions of any external substances. In our current study we obtained the following findings regarding the effects of Koisio technology-modulated solutions on cell growth: First, compared with the lung fibroblast (L929) cell cultures cultured in normal media, the L929 cells cultured in Koisio technology-modulated media grew at approximately 20% higher speed; second, compared with the lung cancer cell cultures (LLC cells) cultured in normal media, the LLC cells cultured in Koisio technology-modulated media grew at approximately 9% lower speed; and third, compared with the telomere lengths of the L929 cells cultured in normal media, the L929 cells cultured in Koisio technology-modulated media had approximately 14% longer telomere length. Collectively, our study has provided the first evidence indicating that Koisio technology-modulated solutions affect differentially the growth of lung fibroblast cell cultures and that of lung cancer cell cultures. The capacity of the Koisio technology-modulated solutions to promote the growth of lung fibroblast cell cultures may result at least partially from its capacity to protect the telomere length.

## Introduction

Cell growth is a critical biological property of cells, which profoundly affects multiple key biological processes such as organ development, tissue repair, and cancer development (1–6). Tissue repair can also be affected by numerous factors such as oxidative stress and NAD^+^ metabolism (7–9). Under injury conditions, enhanced cell growth can lead to accelerated tissue repair (4). In contrast, aberrant and uncontrolled cell growth is a hallmark of cancer cells (5). Cell growth is controlled by multiple factors and highly complex mechanisms (6). Due to the profound significance of cell growth, it is critical to discover novel approaches to enhance normal cell growth and inhibit cancer cell growth.

It has been established that telomere shortening is a hallmark of cellular aging (10, 11). Prevention of telomere shortening is an important strategy for slowing the aging process (10, 11). It is of both scientific and biological significance to search for novel strategies to slow down telomere shortening.

Koisio technology is a novel technology that modulates the properties of water solely by physical approaches without additions of any external substances. A previous study has reported that the ceramics produced by Koisio technology could produce water with increased permeability through aquaporins (12). Our latest study has also found that Koisio technology-modulated media can lead to an approximately 57% increase in intracellular ATP levels. However, it has been unclear if Koisio technology-modulated solutions without additions of any external substances may affect cell growth. Due to the critical biomedical significance of cell growth, in current study we determined the effects of the solutions modulated by Koisio technology on cell growth of both lung fibroblast cell cultures and lung cancer cell cultures. Our study showed that the solutions modulated by Koisio technology produced differential effects on the cell growth of these two types of cells.

## Materials and Methods

### Cell Cultures

Mouse lung fibroblast (L929) cell lines (13, 14) were purchased from Chinese Academy of Sciences Cell Bank. Lung cancer (LLC) cell lines were purchased from American Type Culture Collection (CRL-1642^™^). The cells were grown in DMEM (SH30243.01, Hyclone) supplemented with 10%FBS (100-500, GEMINI), 100 U/mL penicillin, and 0.1 mg/mL streptomycin (15140122, Gibco) at 37 °C in humidified 5% CO_2_ atmosphere.

### Modulations of cell culture media by Koisio technology

Modulations of cell culture media were conducted by Koisio technology as reported previously (12).

### CCK8 assay

Cell viability was measured by using Cell Counting Kit-8 (Dojindo Laboratories, Kumamoto, Japan). In brief, cells were seeded in 96-well plates at the density of 5 × 10^3^ per well. After 24 h, CCK8 solution was added into the medium at a dilution of 1:10 and incubated at 37°C for 1–3 h. The absorbance at 450 nm was determined using a microplate reader (Synergy2, BioTek, Winooski, VT, USA).

### Determinations of telomere lengths by qPCR

Cell’s DNA was extracted by using DNA extraction kit (DP1902, Bioteke Inc, Beijing) according to the manufacturer’s protocol. Mean telomere length was measured by Shanghai Biowing Applied Biotechnology CO. LTD, as described previously (15). qPCR was conducted in triplicate, and the reactions included 4 μl of genomic DNA (80 ng), 0.1 μl of telomere primer (10 μM) (forwards: CGGTTTGTTTGGGTTTGGGTTTGGGTTTGGGTTTGGGTT; reverse: (GGCTTGCCTTACCCTTACCCTTACCCTTACCCTTACCCT), 0.1 μl of YH-1 forwards primer (10 μM) (5′-CGCACAGAGTAGTAAG-GAAAGTGAAGTAGGCCGGGC-3′), 1 μl of YH-1 reverse primer (10 μM) (5′-GTGCTGGGATTACAGGCGTGAG-3′), 1 μl of uniprimer2 (VIC-ATGGACAGTGAGATCTGTCCAT-BHQ1CGCACAGAGTAGTAAG), and 10 μl NovoStart^®^ SYBR qPCR SuperMix, in a final reaction volume of 20 μl.

For PCR product testing, we adopted universal molecular beacon technology and the telomere amplifications were detected using SYBR green dye. All PCRs were carried out on a 7500 Real-Time PCR System, and amplification was conducted as follows: Stage 1:5 min at 95°C, Stage 2:30 cycles (telomere and internal gene reaction) of 15 sec at 95°C, Stage 3:1 min at 50°C, Stage 4:45 sec at 72°C, and fluorescence signals were collected at 50°C. The reactions were carried out in a 96-well plate. Software v2.3 was used for analysis. The telomere length for each sample was determined using the telomere to high-copy gene ratio (T/H ratio) by calculating the ΔCt [Ct(telomere)/Ct (high-copy gene)]. The T/H ratio for each sample (x) was normalized to the mean T/S ratio (test sample/standard sample) of the reference sample [2^−(ΔCtx-2ΔCtr)^=2^−ΔΔCt^], which was also used for the standard curve as a reference sample and as a validation sample. In every run, two reference samples were included to validate each reaction. The experiment was considered reliable if the T/H ratio of the control sample ranged within the 95% variation interval (0.95–1.05).

### Statistical analyses

All data are presented as mean + SEM. Data were analyzed by student *t*-test. *P* values less than 0.05 were considered statistically significant.

## Results

### Lung fibroblast (L929) cell cultures cultured in Koisio technology-modulated DMEM media (KM) showed significantly faster growth speed compared with the L929 cells cultured in regular DMEM media

We determined the effects of KM on the growth of L929 cells – a lung fibroblast cells. Compared with the L929 cells cultured in regular DMEM media, the L929 cells cultured in KM grew at approximately 20% higher speed (Fig. 1).

**Fig.1.**
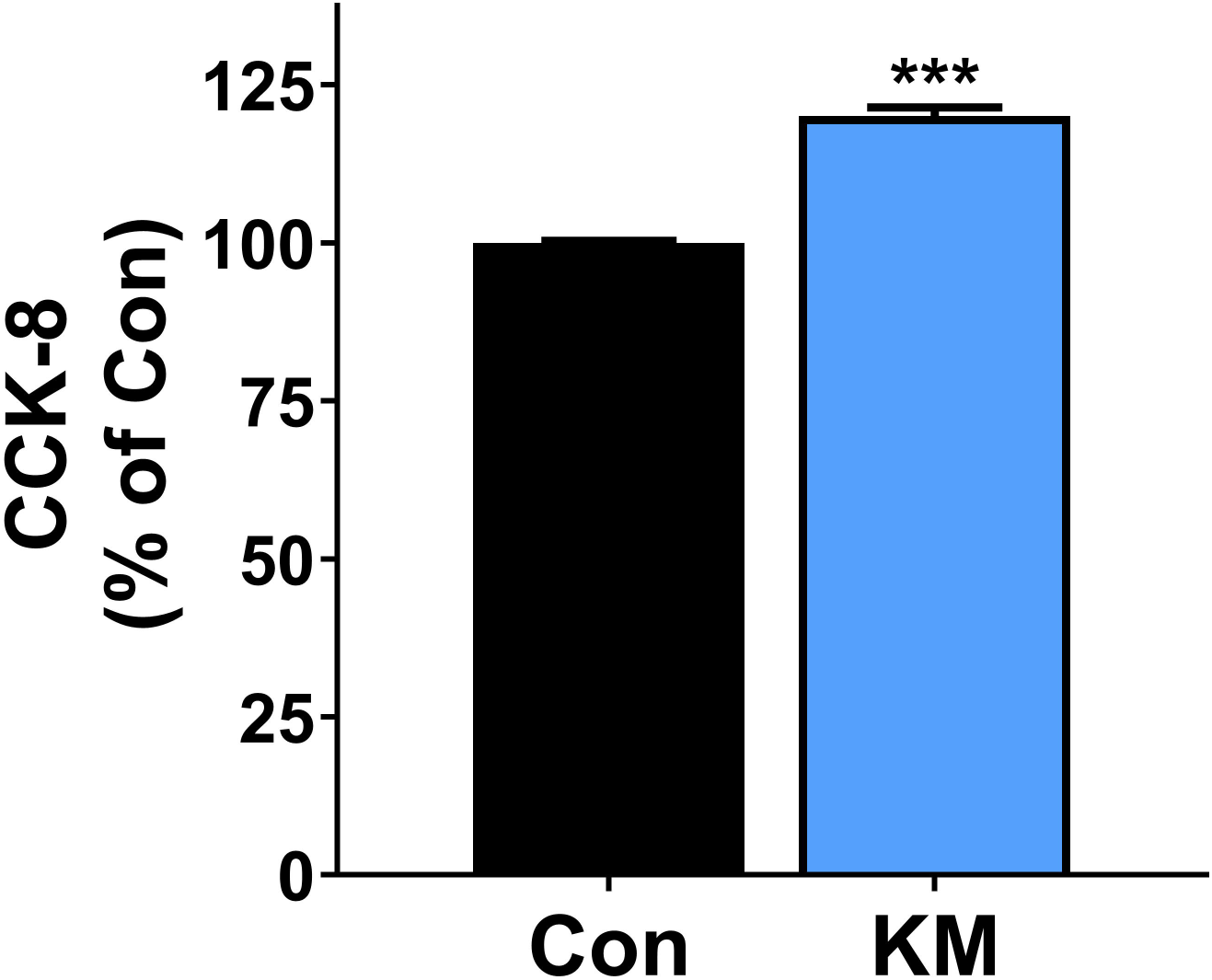
The lung fibroblast (L929 cells) cell cultures cultured in Koisio technology-modulated DMEM media (KM) showed significantly faster growth speed compared with the L929 cells cultured in regular DMEM cell culture media. Compared with the L929 cells cultured in normal cell culture media, the L929 cells cultured in KM grew at approximately 20% higher speed. From Passage 2, a part of the LLC cells grew in the KM. CK assay were conducted when the cell growth reached Passage 10 – 16. The results were collected from five independent CCK assays on five passages of cells. N = 30, *P* < 0.05.

### The lung cancer (LLC) cell cultures cultured in KM showed significantly slower growth speed compared with the lung cancer cell cultures cultured in regular DMEM media

We also determined the effects of KM on the growth of LLC cells – a lung cancer cell line. Compared with the LLC cells cultured in regular DMEM media, the LLC cells cultured in KM did not grow at faster speed, instead, the cells grew at approximately 9% slower speed (Fig. 2).

**Fig.2.**
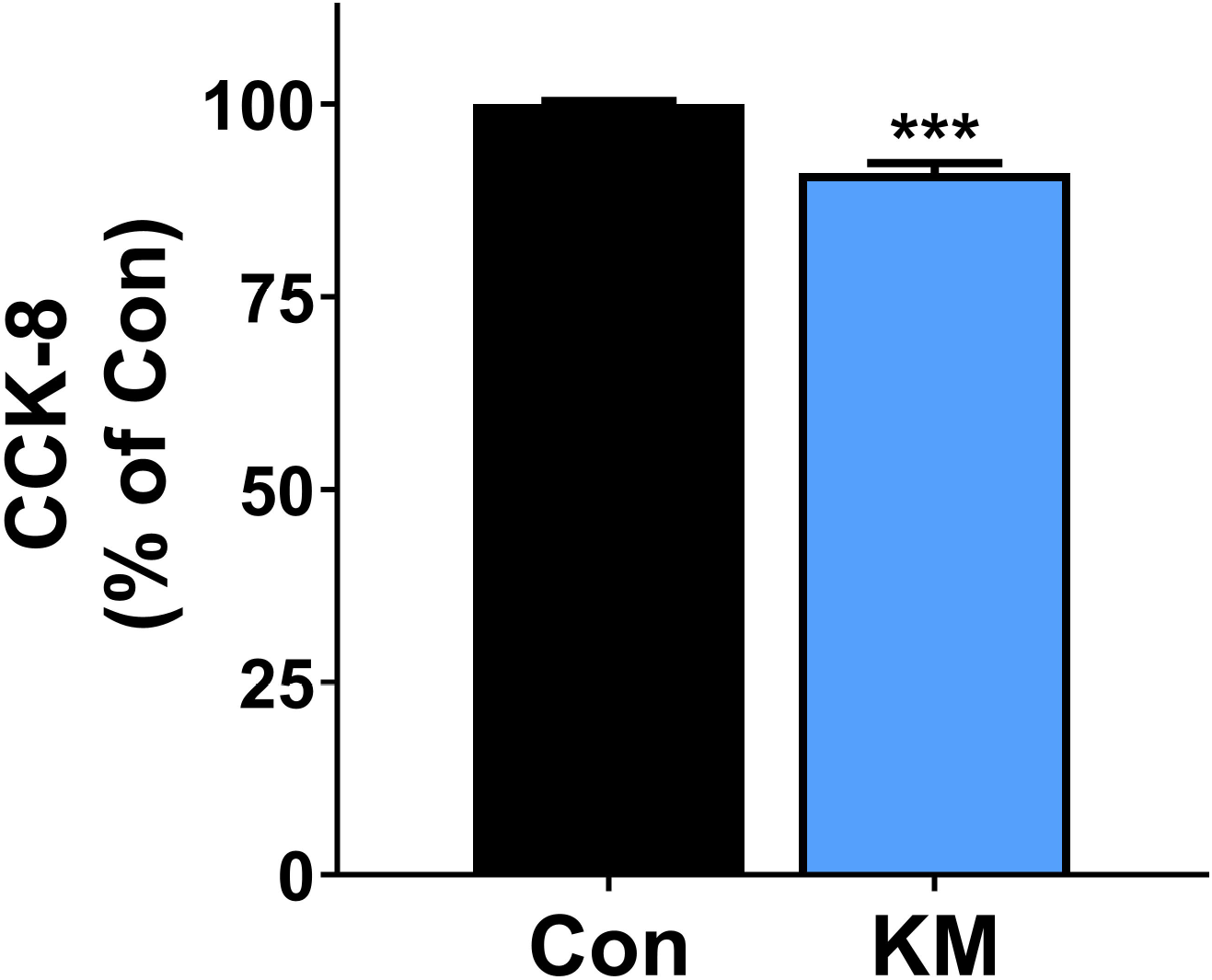
The lung cancer cell cultures (LLC cells) cultured in Koisio technology-modulated DMEM media (KM) showed slower growth speed compared with the LLC cells cultured in regular DMEM cell culture media. Compared with the LLC cells cultured in normal media, the LLC cells cultured in KM grew at approximately 9% lower speed. From Passage 2, a part of the LLC cells grew in the KM. CCK assay were conducted when the cells reached Passage 10 – 16. The results were collected from the five independent CCK assays on five passages of cells. N = 30, *P* < 0.05.

### L929 cells cultured in KM had significantly longer telomere lengths compared with the L929 cells cultured in regular DMEM media

Since telomere shortening is a hallmark of cellular aging, we determined the effects of KM on the telomere length of L929 cells. Compared with the L929 cells cultured in regular DMEM media, the L929 cell cultures cultured in KM had approximately 14% longer telomere length (Fig. 3).

**Fig. 3.**
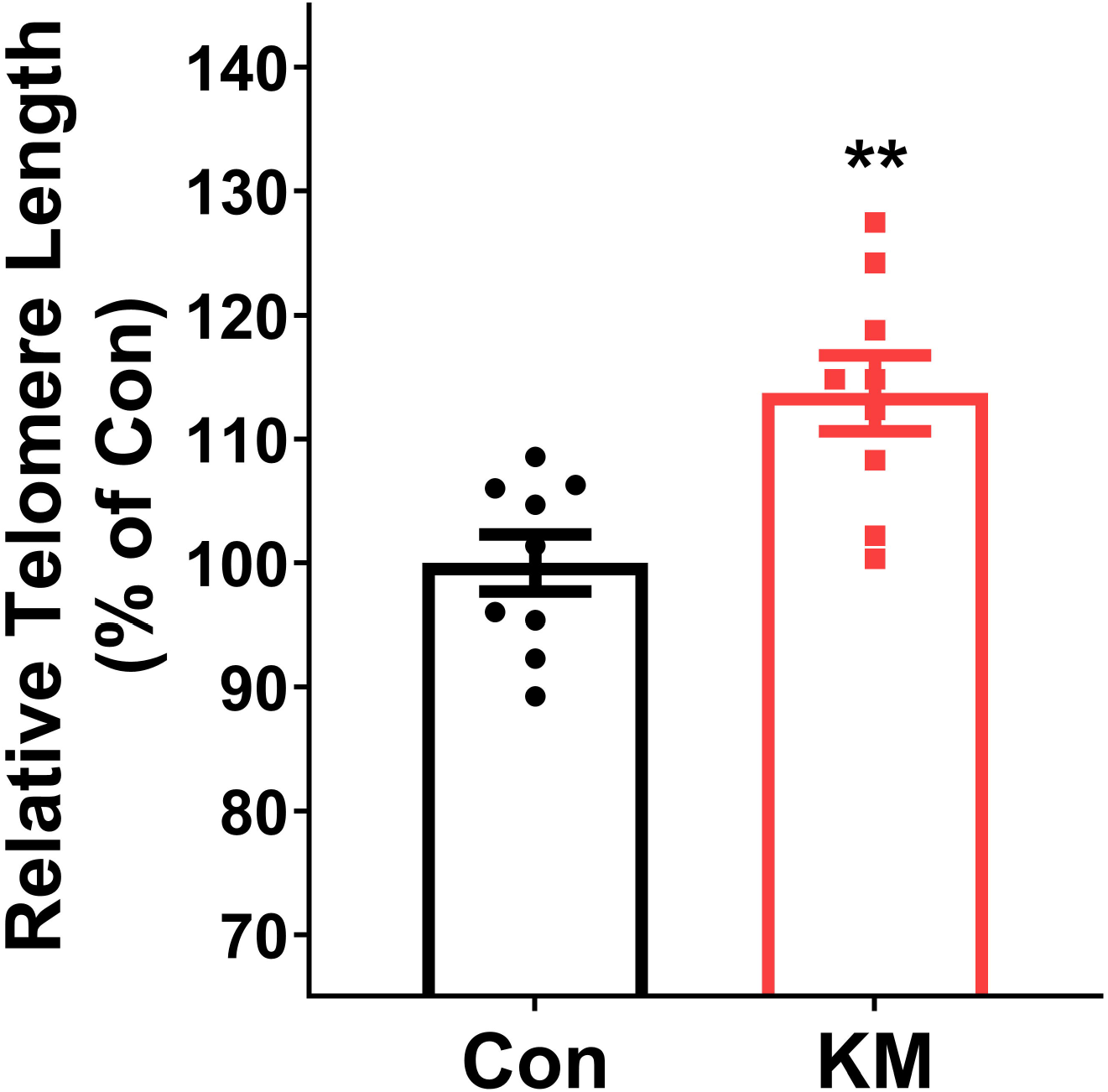
The lung fibroblast (L929 cells) cell cultures cultured in Koisio technology-modulated DMEM media (KM) had significantly longer telomere lengths compared with the L929 cells cultured in regular DMEM cell culture media. Compared with the telomere lengths of the L929 cells cultured in normal cell culture media, the L929 cells cultured in KM had approximately 15% longer telomere length. From Passage 2, a part of the L929 cells grew in KM. Assays of telomere lengths of the cells were conducted when the cells reached Passage 17 – 19. The results were collected from three independent assays on three passages of cells. N = 9, *P* < 0.01.

## Discussion

The major findings of our study include: First, compared with the lung fibroblast (L929 cells) cell cultures cultured in normal media, the lung fibroblast cell cultures cultured in KM grew at approximately 20% higher speed; second, compared with the lung cancer cell cultures (LLC cells) cultured in normal media, the LLC cells cultured in KM grew at approximately 9% lower speed; and third, compared with the telomere lengths of the L929 cells cultured in normal media, the L929 cells cultured in KM had approximately 14% longer telomere length.

The growth rate of normal cells is of great biomedical value. A relatively high growth rate of normal cells can lead to higher speed of tissue repair if an organ or tissue is injured. Our study has indicated that the lung fibroblast cell cultures cultured in KM grew at approximately 20% higher speed, suggesting that Koisio technology-based solutions may enhance the growth rate of normal cells as well as the tissue repair speed.

Cancer cells are cells with aberrant and uncontrolled growth. Therefore, it is critical to determine if KM may also accelerate the growth rate of cancer cells. Our study has found that compared with the LLC cells cultured in normal cell culture media, the LLC cell cultures cultured in KM did not grow at a higher speed, in instead, the cells grew at approximately 9% slower speed.

Collectively, our study has suggested that Koisio technology-modulated media affect differentially the growth rate of normal lung fibroblast cell cultures and lung cancer cell cultures. However, the mechanisms underlying the differential effects of KM on the growth rate of normal lung cells and lung cancer cells remain unclear. It is warranted to conduct future studies to investigate the mechanisms underlying these differential effects.

Telemere shortening is a major index of *in vitro* aging of cells (10, 11). Prevention of telemere shortening is an important strategy for slowing down cellular aging (10, 11). Our study has shown that compared with the telomere lengths of the L929 cells cultured in normal media, the L929 cells cultured in KM had approximately 14% longer telomere length. This finding has suggested that KM may enhance the growth speed of L929 cells at least partially by slowing down the shortening of telemere length of the cells.

